# Modeling Cellular Resource Allocation Reveals Low Phenotypic Plasticity of C_4_ Plants and Infers Environments of C_4_ Photosynthesis Evolution

**DOI:** 10.1101/371096

**Authors:** Esther M. Sundermann, Martin J. Lercher, David Heckmann

**Author notes:** To whom correspondence should be addressed; David Heckmann.

## Abstract

- The regulation of resource allocation in biological systems observed today is the cumulative result of natural selection in ancestral and recent environments. To what extent are observed resource allocation patterns in different photosynthetic types optimally adapted to current conditions, and to what extend do they reflect ancestral environments? Here, we explore these questions for C_3_, C_4_, and C_3_-C_4_ intermediate plants of the model genus *Flaveria*.
- We developed a detailed mathematical model of carbon fixation, which accounts for various environmental parameters and for energy and nitrogen partitioning across photosynthetic components. This allows us to assess environment-dependent plant physiology and performance as a function of resource allocation patterns.
- To achieve maximal CO_2_ fixation rates under growth conditions differing from those experienced during their evolution, C_4_ species need to re-allocate significantly more nitrogen between photosynthetic components than their C_3_ relatives. As this is linked to a limited phenotypic plasticity, observed resource distributions in C_4_ plants still reflect optimality in ancestral environments, allowing their quantitative inference.
- Our work allows us to quantify environmental effects on resource allocation and performance of photosynthetic organisms. This understanding paves the way for interpreting present photosynthetic physiology in the light of evolutionary history.

## Introduction

Metabolic efficiency is an important determinant of organismal fitness (Ibarra *et al.*, 2002; Heckmann *et al.*, 2013). Major constraints on metabolic fluxes can arise from scarcity of chemical compounds, e.g., nitrogen necessary to produce enzymes (Baudouin-Cornu *et al.*, 2001), or from the limited solvent capacity of cellular compartments (Atkinson, 1969; Beg *et al.*, 2007). To ensure optimal metabolic efficiency, gene regulation has to balance available resources appropriately. Modern methods of modeling metabolism rely strongly on the assumption of metabolic optimality under physico-chemical constraints (Oberhardt *et al.*, 2009; de Oliveira Dal’Molin *et al.*, 2010; Dourado *et al.*, 2017). Accordingly, resource allocation and its constraints are under intense investigation, although these studies are mostly restricted to unicellular organisms. However, the metabolic efficiency of a given metabolic system is not static, but depends on the environment. Thus, uncertainties about the environmental properties that an organism has adapted to remain a major obstacle in the application of these methods. Autotrophic systems, such as plant leaves, are ideal to study the interaction of the environment and resource allocation, as the diversity of nutrient sources is much lower than for heterotrophs, which results in a reduced complexity of the space of possible environments. Furthermore, the effect of environmental factors on plant performance, e.g., the rate of CO_2_ assimilation, have been studied intensively (von Caemmerer, 2000). In particular, C_3_ and C_4_ photosynthesis represent complementary gene expression and resource allocation patterns that result in high fitness in specific ecological niches.

In all plants, the fixation of carbon from CO_2_ is catalyzed by the enzyme ribulose-1,5-bisphosphate carboxylase/oxygenase (Rubisco) as part of the Calvin-Benson cycle. Rubisco also shows an affinity for O_2_, resulting in a toxic by-product, which needs to be recycled by the photorespiratory pathway and causes a significant loss of carbon and energy (Maurino & Peterhansel, 2010). Rubisco is an important resource sink in the leaf proteome of plants: it utilizes up to 30% of leaf nitrogen and up to 65% of total soluble protein (Ellis, 1979; Makino *et al.*, 2003). While C_3_ plants operate the Calvin-Benson cycle in their mesophyll cells to fix carbon, C_4_ plants express it in the bundle sheath cells and use phospho*enol*pyruvate (PEP) carboxylase (PEPC) for the initial fixation of carbon. The resulting C_4_ acids are eventually decarboxylated in the bundle sheath cells, creating a local high-CO_2_ environment around Rubisco that suppresses photorespiration. The C_4_ cycle is completed by the regeneration of PEP by pyruvate, phosphate dikinase (PPDK).

Compared to C_3_ photosynthesis, C_4_ metabolism requires additional nitrogen to produce the C_4_ enzymes; this additional investment is counteracted by reduced Rubisco requirements due to the concentration of CO_2_ around Rubisco (Sage, 2004). The energy requirements of C_4_ metabolism also differ from those of the C_3_ pathway (Munekage & Taniguchi, 2016), as further ATP is needed for the regeneration of PEP, while ATP and NADPH requirements of the photorespiratory pathway are reduced. The metabolic efficiencies of the C_3_ and C_4_ system depend strongly on the environment. To achieve optimal metabolic efficiency, plants have to coordinate gene expression of the Calvin-Benson cycle, C_4_ cycle, photorespiration, and light reactions in a complex response to the availability of light energy and nitrogen, as well as factors that influence the rate of photorespiration. The diversity of photosynthetic resource allocation patterns is emphasized by the existence of C_3_-C_4_ intermediate photosynthesis in some plants, where features of the archetypical C_4_ syndrome are only partially expressed. The *Flaveria* genus contains closely related plants of C_3_, C_4_, and C_3_-C_4_ intermediate types, making it an ideal system to study the interaction between resource allocation and environment in photosynthesis.

The selection pressures caused by environmental factors over evolutionary time scales are expected to lead to corresponding adaptations of gene regulation. In contrast, environmental variation on the time scale of individual generations may select for regulatory programs that adjust plant metabolism to the environment they currently face, a process called phenotypic plasticity. Reviewing the occurrence of phenotypic plasticity in C_3_ and C_4_ plants, Sage and McKown (2006) concluded that C_4_ plants show inherent constraints that prevent the acclimation to environmental changes. Although the occurrence of phenotypic plasticity in plants is intensively studied, the plasticity in terms of resource allocation is not fully understood. In particular, it is not clear whether the phenotypic plasticity of different plant lineages is sufficient to acclimate optimally to the current environment; instead, many plants might still allocate at least parts of their resources in patterns that were optimal in the environments that dominated their recent evolutionary history.

The areas where C_4_ dicotyledonous plants are assumed to have evolved are regions of low latitude showing combinations of heat, drought, and salinity (Sage, 2004). For *Flaveria*, analyses that combine phylogenetic context and environmental information point toward an evolutionary origin in open habitats with high temperatures (Powell, 1978; Sage, 2004; McKown *et al.*, 2005). The last common C_3_ ancestor of the current *Flaveria* species lived 2–3 million years ago (Christin *et al.*, 2011), when CO_2_ levels were significantly lower than the current, postindustrial level (Sage & Cowling, 1999; Gerhart & Ward, 2010). In summary, *Flaveria* species likely faced high light intensities, high temperature, and low atmospheric CO_2_ level during their recent evolutionary history.

Here, we aim for a detailed understanding of the interplay between resource allocation and current and past evolutionary environments in plant physiology, examining C_3_, C_4_, and C_3_-C_4_ intermediate photosynthesis. To achieve this goal, we developed a mathematical model for these photosynthetic types that integrates knowledge on resource costs and relevant environmental factors. Using this model, we seek to understand (1) to what extent resource allocation is phenotypically plastic and to what extent it appears adapted to an environment the plants were facing during their evolutionary history; and (2) if resource allocation patterns can be used to identify unique environments of optimal adaption.

## Results

### Predicting resource allocation and fitness across environments and photosynthetic types: a mathematical model

The standard method to model the light-and enzyme-limited CO_2_ assimilation rate of C_3_, C_4_, and C_3_-C_4_ intermediate plants is based on the mechanistic biochemical models of Berry and Farquhar (1978), Farquhar *et al.* (1980), and von Caemmerer (1989; 2000). With great success, these models predict the CO_2_ assimilation rate considering enzymatic activities and various environmental parameters, including mesophyll CO_2_ level and light intensities. In many ecosystems, the most limiting resource for plant growth is nitrogen (Malhi *et al.*, 2001; Vance, 2001). The increased nitrogen-use efficiency of C_4_ species compared to C_3_ relatives indicates that nitrogen availability may have played a major role in C_4_ evolution (Vogan & Sage, 2011). However, existing model implementations predict CO_2_ assimilation rates from known or estimated enzyme activities and electron transport capacity. Thus, these models do not allow to assess the effects of nitrogen investment into different classes of proteins—including enzymes and components of the electron transport chain—on the CO_2_ assimilation rate of a given photosynthetic type in a specific environment.

Here, we present a nitrogen-dependent light-and enzyme-limited model for the steady-state CO_2_ assimilation rate (Fig. 1). The model describes C_3_, C_4_, and all intermediate photosynthetic types depending on its parameterization, including the nitrogen investment into its different components (see Heckmann *et al.* (2013) for details and Supporting Information for our parameterization). We modified the light-and enzyme-limited C_3_-C_4_ models developed by von Caemmerer (2000) and added a fixed budget of nitrogen constraining the total abundance of photosynthetic proteins. Furthermore, we extended the existing models by explicitly modeling the ATP and NADPH production of the linear and cyclic electron transport (LET and CET, respectively). Thus, a photosynthetic nitrogen budget is distributed across the enzymes of the Calvin-Benson cycle in the mesophyll and bundle sheath cell, the C_4_ cycle, and the proteins of the linear and cyclic electron transport in the thylakoid membranes. Combining this model with the temperature dependency of the photosynthetic apparatus (Massad *et al.*, 2007) results in a detailed model of photosynthesis that incorporates leaf nitrogen level, light intensity, mesophyll CO_2_ and O_2_ levels, as well as the effects of temperature (see Methods for details).

**Figure 1.**
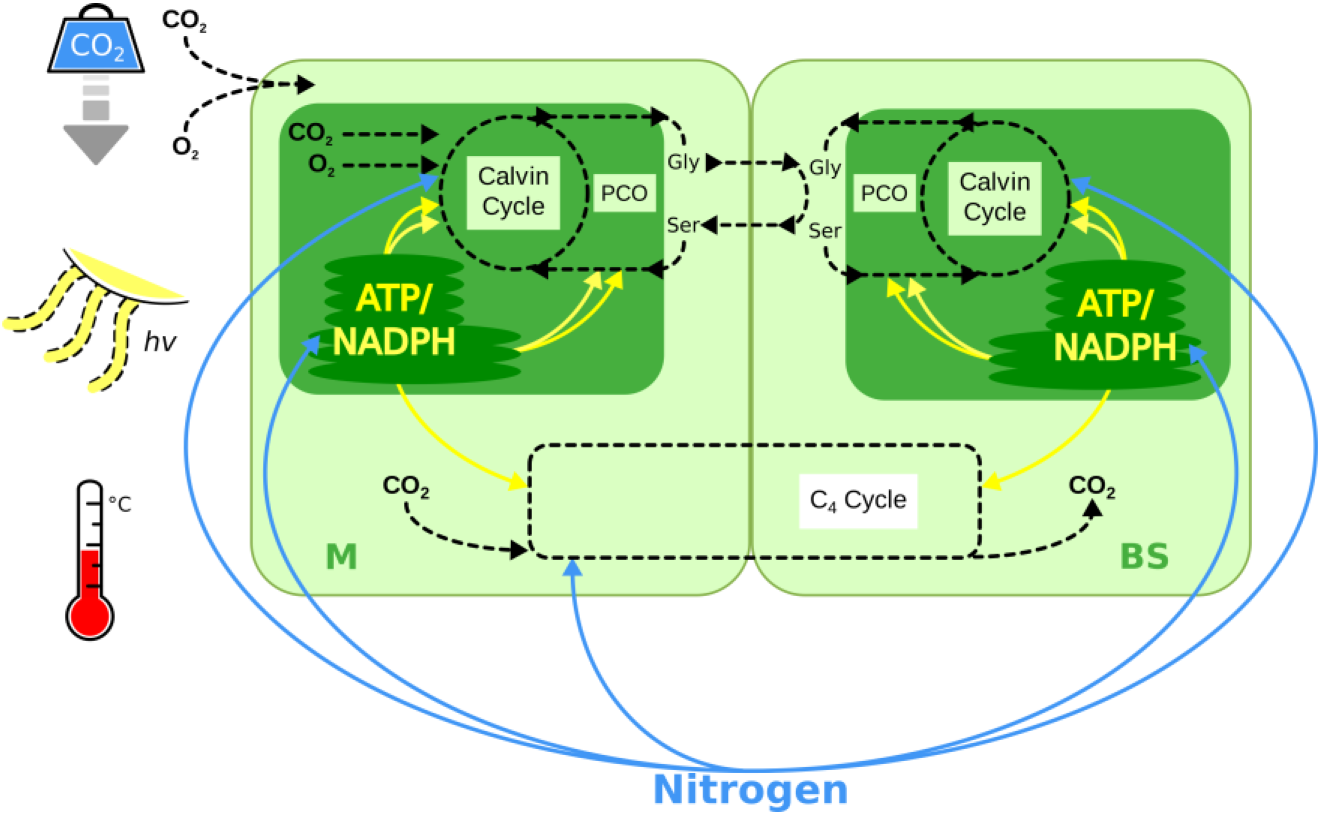
An overview of the nitrogen-dependent light-and enzyme-limited model. CO_2_ entering the mesophyll cell (M) can be fixed by Rubisco (C_3_ and intermediates) or PEPC (C_4_ and intermediates); The C_4_ cycle then shuttles CO_2_ fixed by PEPC to the bundle sheath cell (BS) and releases it, allowing it to be re-fixed by Rubisco. The fixation of O_2_ by Rubisco leads to photorespiration (PCO). Blue arrows indicate the nitrogen allocation and yellow arrows represent the energy allocation considered in the model.

In order to understand physiological data in the context of adaptive environments, we aim to find optimal resource allocation in a given environment. To this end, we assume that resource allocation has been optimized by natural selection to maximize the net CO_2_ assimilation rate (Zhu *et al.*, 2007; Gerhart & Ward, 2010; Vogan & Sage, 2012). We developed a robust optimization pipeline that reliably finds optimal resource allocation dependent on environments and photosynthetic types (see Methods for details). In previous work, optimality assumptions were successfully used in a variety of plant systems biology contexts; examples are candidate identification of photosynthetic engineering targets (Zhu *et al.*, 2007), explanation of the coordination of C_3_ photosynthesis (Friend, 1991; Maire *et al.*, 2012), the exploration of evolutionary trajectories of C_4_ photosynthesis (Heckmann *et al.*, 2013) and of inter-cellular pathways in C_2_ plants (Mallmann *et al.*, 2014), and the prediction of dynamic proteome allocation in cyanobacteria (Reimers *et al.*, 2017). We use optimality of CO_2_ fixation rate to determine (1) the optimal partitioning of NADPH between the Calvin-Benson cycle and the photorespiratory pathway, (2) the optimal partitioning of ATP across the Calvin-Benson cycle, photorespiratory pathway, and the C_4_ cycle (if relevant), (3) the optimal proportion of LET and CET, and (4) the relative investment of nitrogen into Rubisco, the C_4_ cycle enzymes, and the proteins of the light-dependent reactions (see Methods). For a specific photosynthetic type, the optimization procedure estimates the resource allocation that is optimally adapted to a given environment. Note that at the point of optimal resource allocation, the light-and enzyme-limited CO_2_ assimilation rates are equal, as otherwise resources could be shifted from the non-limiting to the limiting sector.

### Optimal resource allocation in the evolutionarily relevant environment explains physiological data and outperforms models based on the growth environment in C_4_ plants

Do photosynthetic types exhibit differences in phenotypic plasticity, *i.e.*, do they differ in their ability to adjust their photosynthetic resource allocation to optimally fit the environment in which they were grown? Or is resource investment static and reflects past environments in which the plants’ ancestors evolved? To compare these competing hypotheses, we predict physiological data of plants that are either optimally adapted to the experimental growth conditions used in the respective studies (‘growth scenario’) or to the environments in which they likely evolved (‘evolutionary scenario’). This *in silico* experiment also serves as validation for our modeling framework; if the parameterization for *Flaveria* and our optimality assumptions are correct, we would expect the model to explain physiological responses in one of the two or an intermediate scenario. Based on the suggested environment of C_4_ evolution in *Flaveria* (Powell, 1978; Sage, 2004; McKown et al., 2005), the evolutionary environment is defined as having 1750 μmol quanta m^−2^ s^−1^ light intensity, 30°C temperature, 150 μbar mesophyll CO_2_, and 200 mbar mesophyll O_2_.

Vogan and Sage (2012) measured the net CO_2_ assimilation rate as a function of intercellular CO_2_ concentration (A-C_i_ curve) for *Flaveria robusta* (C_3_), *F. ramosissima* (C_3_-C_4_), and *F. bidentis* (C_4_). In this experiment, plants were grown at light intensities of 560 μmol quanta m^−2^ s^−1^, 37°C at daytime, current atmospheric O_2_ concentration and current or low atmospheric CO_2_ concentrations. However, CO_2_ assimilation curves calculated from a model parameterized for optimal CO_2_ assimilation in these growth conditions are qualitatively different from the experimental curves (Fig. 2a; Supporting Information Figs. S2–S4). In contrast, the modeled curves based on a model optimally adapted to the evolutionary scenario are qualitatively consistent with the measured curves; this difference is especially pronounced in the case of the C_4_ plant *F. bidentis*.

**Figure 2.**
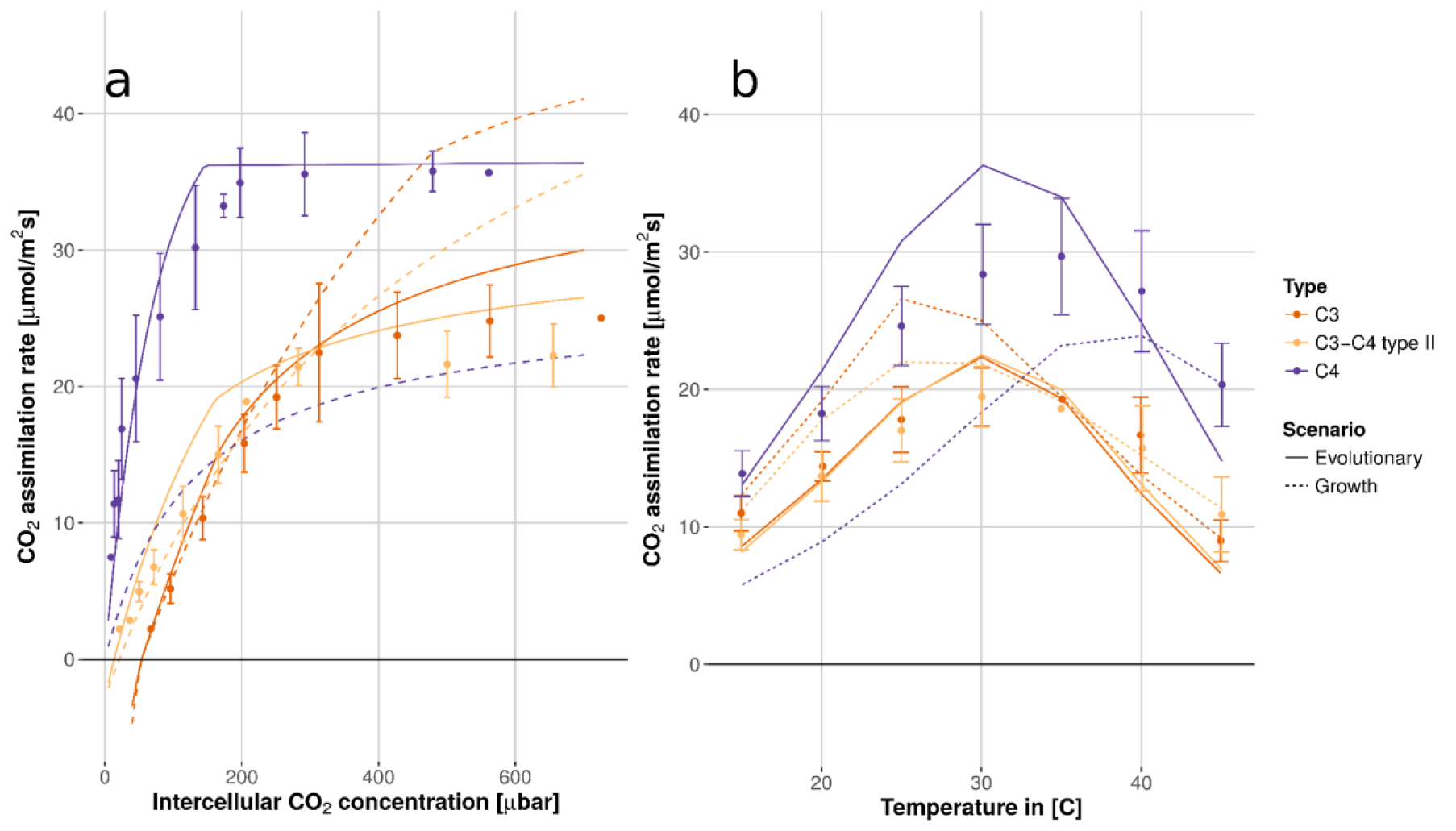
Model results based on optimality in the evolutionary scenario (solid lines) describe the measured data (dots ± SE) better than the model assuming optimal adaptation to the growth conditions (dashed lines) for *F. robusta* (C_3_), *F. ramosissima* (C_3_-C_4_), and *F. bidentis* (C_4_) grown at the current CO_2_ level (data from Vogan and Sage (2012)). (a) The net CO_2_ assimilation rate as a function of intercellular CO_2_ concentration measured at 30°C. (b) The net CO_2_ assimilation rate as a function of temperature.

In the same study, Vogan and Sage (2012) measured the CO_2_ assimilation rate for temperatures between 15°C and 45°C (A-Temperature curve; Fig. 2b; Supporting Information Fig. S5). The results assuming an optimal allocation under the evolutionary scenario agree qualitatively with the measured data, again in contrast to the values predicted from a model optimally adapted to the growth environment. Note that none of the species in this data set were used to obtain the temperature response curves used in the model (see Methods).

In an independent experiment, Vogan and Sage (2011) measured the dependence of CO_2_ assimilation rate on leaf nitrogen levels in C_3_, C_3_-C_4_ intermediate, C_4_-like, and C_4_ *Flaveria* species (Fig. 3). The plants were grown at 554 μmol quanta m^−2^ s^−1^ light intensity, 30°C at daytime, at current atmospheric CO_2_ and O_2_ concentrations. Again, the model results assuming optimal resource allocation in the evolutionary scenario are consistent with the measured data and outperform the results based on optimality in the growth scenario for C_3_-C_4_ intermediate, C_4_-like, and C_4_ plants.

**Figure 3.**
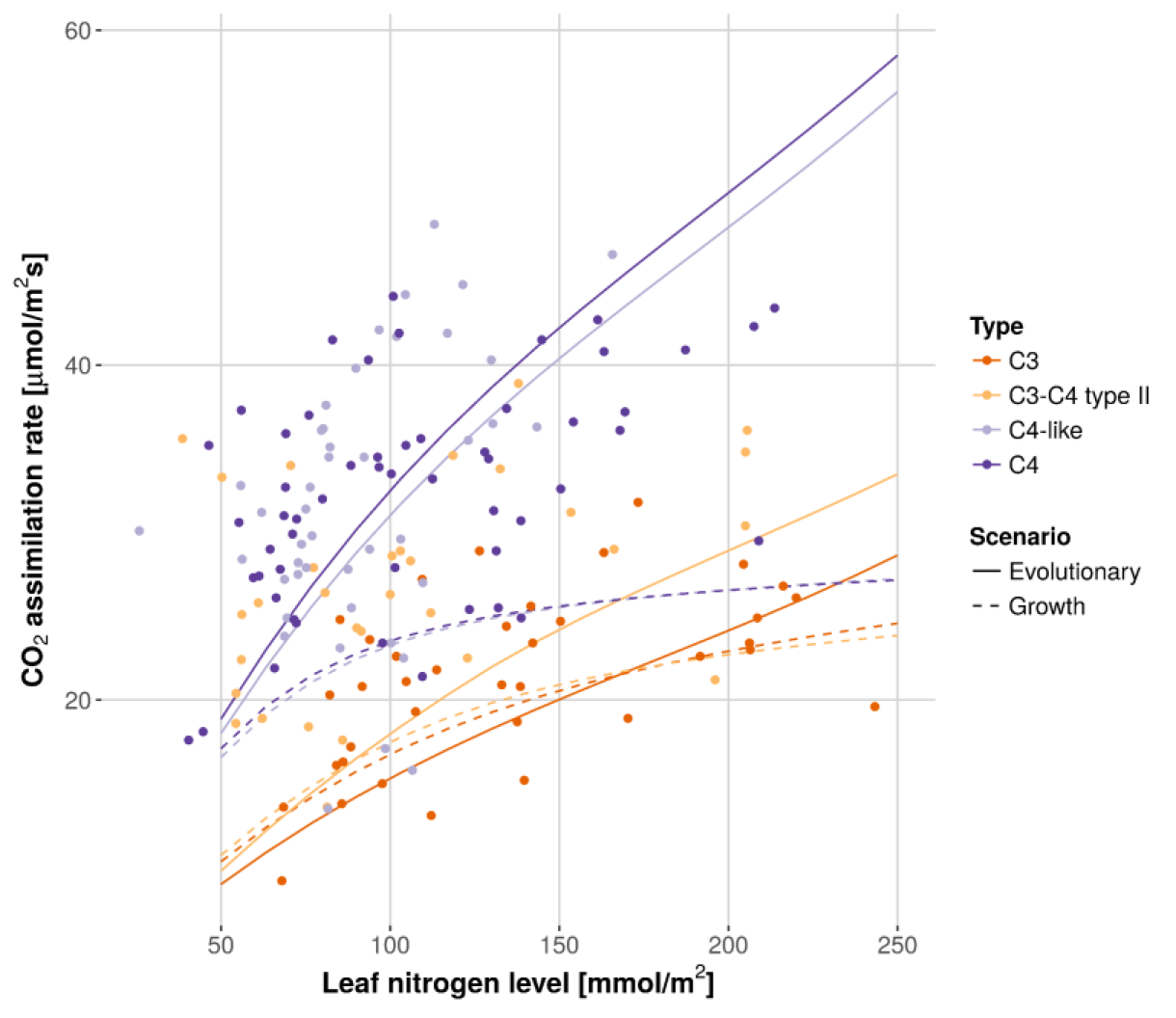
The dependence of the CO_2_ assimilation rate on leaf nitrogen levels for various *Flaveria* species is consistent with model results based on optimality in the evolutionary scenario (solid lines). For C_3_-C_4_ intermediate, C_4_-like, and C_4_ these results outperform the ones assuming optimal phenotypic adaptation to the growth conditions (dashed lines). The modeled species are *F. pringlei* (C_3_)*, F. floridana* (C_3_-C_4_)*, F. palmeri* (C_4_-like)*, and F. bidentis* (C_4_) (data from Vogan and Sage (2011)).

We quantified the disagreement between measured curves and predicted results through the residual sum of squares (Table 1). In C_4_ and C_4_-like plants, the evolutionary scenario predicts all measured curves better than the growth scenario, except for the A-Temperature curve for C_4_ plants grown at low CO_2_ concentration. Jointly considering all measured curves in Figs. 2 and 3 as well as Supporting Information Figs. S2–S5 (Vogan & Sage, 2011; Vogan & Sage, 2012), we find that for the C_4_ and C_4_-like species, squared residuals for the evolutionary scenario are statistically significantly smaller than for the growth scenario (C_4_: *P* = 6.0×10^−8^; C_4_-like: *P* = 0.007; Wilcoxon rank sum tests). This finding indicates that observed resource allocation patterns in C_4_ and C_4_-like plants reflect past environments relevant during evolution more than the environment in which the assayed plants were grown. Conversely, and as expected from Table 1, the observed differences between predictions from the evolutionary and growth scenario are not statistically significant for the C_3_ and the C_3_-C_4_ intermediate species (C_3_: *P* = 0.35; C_3_-C_4_: *P* = 0.55).

**Table 1.** In C_4_ and C_4_-like plants, the evolutionary scenario shows significantly smaller residual sum of squares compared to the growth scenario. The residual sum of squares for the evolutionary and growth scenario, each photosynthetic type, and all measured curves of Vogan and Sage (2011) and Vogan and Sage (2012) are presented.

|  |  | C<sub>3</sub> | C<sub>3</sub>-C<sub>4</sub><br>intermediate | C<sub>4</sub>-like | C<sub>4</sub> |
| --- | --- | --- | --- | --- | --- |
| Evolutionary scenario | Fig. 2a | 58.1 | 77.1 |  | 93.4 |
|  | Fig. 2b | 823.7 | 524.6 |  | 155.6 |
|  | Fig. 3 | 549.2 | 1554.3 | 1443.5 | 834.9 |
|  | Supporting Information Fig. S2 | 616.3 | 299.1 |  | 136.9 |
|  | Supporting Information Fig. S3 | 39.9 | 40.4 |  | 166.0 |
|  | Supporting Information Fig. S4 | 137.0 | 85.6 |  | 286.4 |
|  | Supporting Information Fig. S5 | 14.5 | 93.9 |  | 275.5 |
| Growth scenario | Fig. 2a | 602.2 | 238.2 |  | 2454.5 |
|  | Fig. 2b | 755.2 | 306.1 |  | 340.3 |
|  | Fig. 3 | 386.5 | 2122.2 | 3052.2 | 1873.2 |
|  | Supporting Information Fig. S2 | 252.7 | 84.9 |  | 433.9 |
|  | Supporting Information Fig. S3 | 140.84 | 50.7 |  | 436.1 |
|  | Supporting Information Fig. S4 | 97.8 | 38.8 |  | 460.0 |
|  | Supporting Information Fig. S5 | 13.4 | 53.9 |  | 142.8 |

Dwyer *et al.* (2007) performed detailed experiments on the photosynthetic resource allocation and performance of the C_4_ species *F. bidentis*. This data allows us to compare the predicted nitrogen investment into the three major photosynthetic components—Rubisco, C_4_ cycle, and electron transport chain—as well as the corresponding CO_2_ assimilation rate to experimentally observed resource allocation patterns. The plants were grown under 25°C or 35°C at daytime, 550 μmol quanta m^−2^ s^−1^, and current atmospheric CO_2_ and O_2_ concentrations. Model predictions of chlorophyll content and the amount of photosystem II agree within a factor of 1.10 to 1.22 with values measured by Dwyer *et al.* (2007) (see Supporting Information Table S7). For plants grown at 25°C, the resource allocation determined under the evolutionary scenario agrees with the measured data within a factor of 0.47 to 1.22 (Fig. 4a); at 35°C, agreement is within a factor of 0.43 to 1.12 (Fig. 4b). In both cases, agreement is much lower for predictions in the growth scenario. We assessed the statistical significance of the superior performance of the evolutionary scenario by comparing the distributions of the squared residuals (expressed as fractions of the experimental means). The resource allocation calculated for the evolutionary scenario outperforms the growth scenario significantly for the data represented in Fig. 4 (*P* = 7.2×10^-5^, Wilcoxon rank sum test).

**Figure 4.**
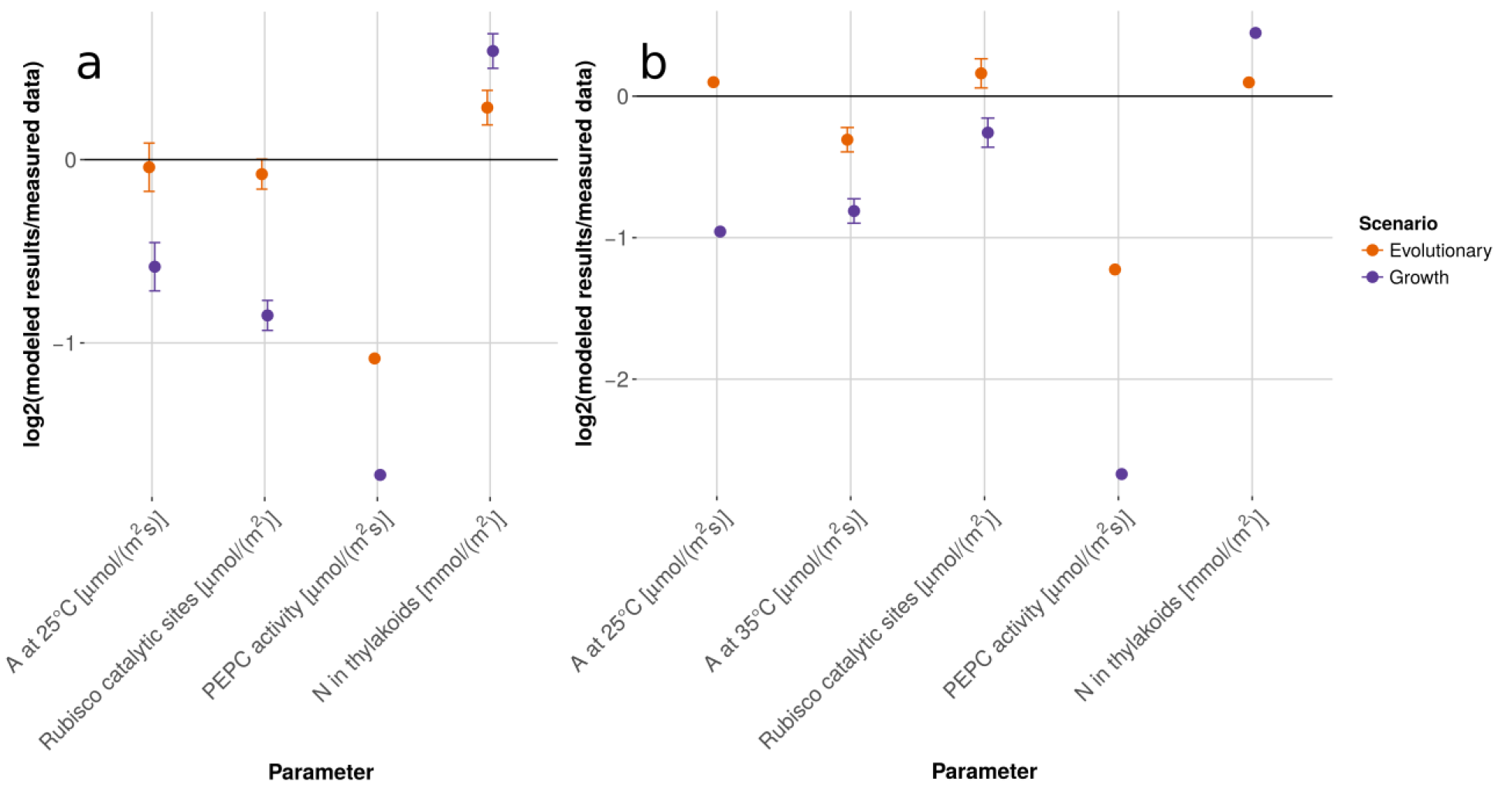
A detailed analysis of resource allocation and physiology in *F. bidentis* (C_4_) shows a good agreement between experimental data (Dwyer *et al.*, 2007) and model results based on the evolutionary scenario (orange dots). Alternative model results assuming optimal phenotypic adaptation to the growth scenario consistently show higher disagreement with the data (purple dots). Values are mean log2(modeled results/measured data) ± SE. (a) Plants grown at 25°C (b) Plants grown at 35°C. *A* = net CO_2_ assimilation rate; *N* = nitrogen.

Although we could obtain the majority of our model parameters from the literature, the relationship of cytochrome f and the maximal electron transport rate of the CET had to be estimated (see Methods). We performed a sensitivity analysis to examine the robustness of the results to changes in the estimated parameters and to uncertainties in values obtained from the literature, focusing on parameters with high uncertainty or major expected effect on model predictions (see Supporting Information Method S5). The predictions based on the evolutionary scenario outperform those based on the growth environment consistently across all parameter sets (Supporting Information Fig. S1).

### The model identifies a unique evolutionary environment for C_4_ photosynthesis in Flaveria

The model optimally adapted to the evolutionary scenario leads to superior predictions of plant performance and resource allocation in C_4_ plants compared to a parameterization optimized for the growth scenario across diverse physiological data sets. The inferior performance of the growth scenario model indicates a lack of phenotypic plasticity of resource allocation in C_4_ plants. This finding points to the possibility that the environment most relevant for recent evolutionary adaptation of a given C_4_ plant could be inferred quantitatively from observations on plant physiology and resource allocation. Thus, to infer a typical evolutionary environment for C_4_ *Flaveria bidentis,* we calculated optimal resource allocation under conditions covering plausible ranges of mesophyll CO_2_ partial pressure, temperature, and light intensities to identify the conditions that best explain the empirical data (Fig. 5). As atmospheric O_2_ concentration remained almost constant for at least the last few million years (Gerhart & Ward, 2010), this environmental parameter is set to a constant value. We use the empirical data of Dwyer *et al.* (2007), as this data set comprises detailed measurements for each nitrogen pool and the resulting CO_2_ assimilation rate, allowing us to quantify the discrepancy between modeled and measured values as the mean squared residuals (expressed as fractions of experimental means).

**Figure 5.**
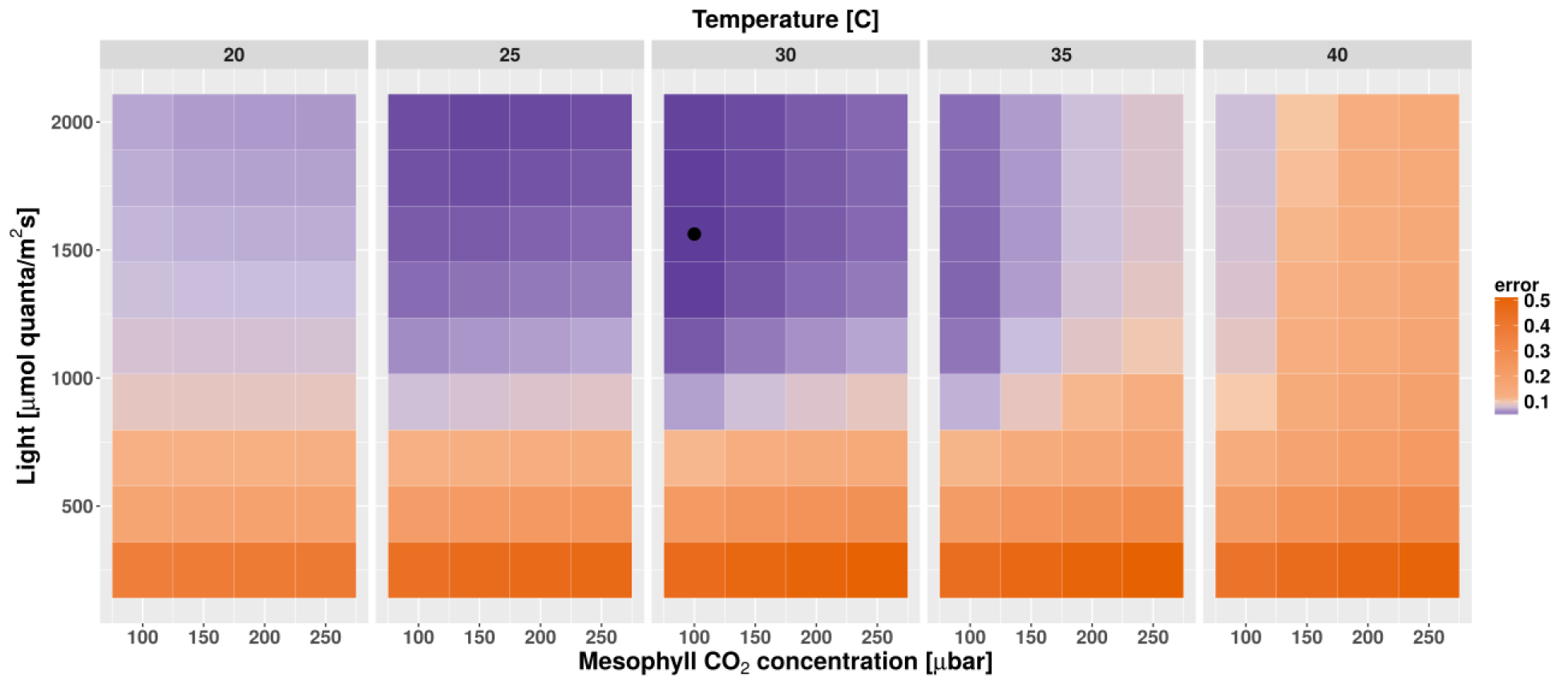
Discrepancy between measured and modeled *F. bidentis* data across diverse environments. The black dot indicates the environment that best explains the experimental data of Dwyer *et al.* (2007). The deviation between model predictions and measurements (‘error’) is defined as the mean of the squared residuals (which are expressed as fractions of experimental means).

We find that the model showing minimal prediction error defines a unique environment (Fig. 5), exhibiting 1562.5 μmol quanta m^−2^ s^−1^ light intensity, 30°C, a mesophyll CO_2_ level of 100 μbar, and an O_2_ level of 200 mbar. As indicated in Fig. 5, the areas in which the model successfully describes the empirical values generally show high light intensities, intermediate to high temperatures, and a trend towards low CO_2_ partial pressures. High light intensities and low CO_2_ levels, as in the evolutionary scenario, favor an increased nitrogen investment into the dark reactions, which goes along with a reduced investment into the electron transport chain. The effect of temperature is of special importance for plants using the C_4_ cycle, as temperature increases PEPC activity drastically and therefore reduces the necessary nitrogen investment into the C_4_ cycle. This allows an increased investment into the electron transport chain and Rubisco, which show reduced activity at elevated temperatures due to thermal instabilities.

Our results indicate that C_4_ *Flaveria* species show a lower degree of photosynthetic phenotypic plasticity than closely related C_3_ species. On a molecular level, this plasticity predominantly requires the re-allocation of nitrogen between the major photosynthetic protein pools. To assess the costs of phenotypic plasticity, we thus quantified the total fraction of nitrogen that needs to be re-allocated between photosynthetic pools to optimally adjust photosynthesis from the evolutionary scenario to a given growth environment (*δ_n_*, see Methods). We find that photosynthetic types that utilize the C_4_ cycle require a consistently higher amount of re-allocation compared to C_3_ plants (*P* = 1.5×10^-5^, sign test, see Supporting Information Table S5). Our results thus reveal a link between required nitrogen re-allocation and limited photosynthetic phenotypic plasticity (see Supporting Information Tables S4–S6), suggesting a possible causal relationship.

## Discussion

Our novel modeling framework allows us to study the interplay between photosynthetic plant performance and resource investment on the molecular level. Comparisons of model predictions with phenotypic and molecular data reveal that C_4_ plants have low phenotypic plasticity in terms of resource allocation. This limited phenotypic plasticity may be explained by the large amount of nitrogen that needs to be re-allocated by C_4_ plants to optimally adapt to a given growth environment (Supporting Information Table S5). The lack of phenotypic plasticity allowed us to make quantitative predictions for the environments that dominated recent evolution of C_4_ photosynthesis in *Flaveria*. Previously, environments relevant for C_4_ photosynthesis evolution have been inferred—mostly qualitatively—based on C_3_-C_4_ habitat comparisons (Powell, 1978; Sage, 2004; McKown *et al.*, 2005) and geophysiological considerations (Christin *et al.*, 2011). Our results are consistent with and refine these earlier estimates.

In contrast to our findings for C_4_ and C_4_-like plants, the performance of the evolutionary and the growth scenario models is similar for C_3_ and C_3_-C_4_ intermediate *Flaveria* species (Table 1; Figs. 2 and 3; Supporting Information Figs. S2–S5). It is conceivable that the lack of superior performance for the evolutionary scenario in C_3_ *Flaveria* species is not a result of higher phenotypic plasticity in these plants, but is due to an inappropriate parameterization of the evolutionary scenario. The environment most relevant for the recent evolution of C_3_ *Flaveria* may be different from the environment used in the simulations, which was chosen based on its relevance for the C_4_ lineages. To explore this possibility, we simulated a wide range of alternative environments, testing if resource allocation optimized for any of these leads to significantly improved model predictions for the data from Vogan and Sage (2012) for C_3_ plants (Supporting Information Figs. S6 and S7). However, none of the environments tested led to a significant improvement. This result is in agreement with habitat studies that show that niches of C_3_ and C_4_ *Flaveria* species overlap (Powell, 1978). A more likely explanation for the similar performance of evolutionary and growth scenario models in C_3_ plants could lie in the small amount of re-allocation C_3_ plants require to transfer adaptively between environments (Supporting Information Tables S4–S6). Our results thus suggest that C_3_ (but not C_4_) plants are phenotypically plastic enough to show some degree of adaptation towards current, postindustrial conditions.

Given the complexity of our physiological model, we needed to make a number of assumptions. We addressed uncertainties in model parameters through a sensitivity analysis, showing that our conclusions are robust against variation in these parameters (Supporting Information Fig. S1). Furthermore, our predictions assume that nitrogen availability in the evolutionary scenario is identical to current nitrogen availability.

Even though we find that the evolutionary scenario leads to superior predictions of physiological responses in C_4_ plants when compared to the growth scenario, the PEPC activity predicted to be optimal in the evolutionary scenario is approximately 55% lower than experimentally observed data (Fig. 4). This discrepancy might in part be explained by the assumption of a fixed average daytime temperature in the simulations. Temperature variation strongly affects the PEPC activity; lower temperatures in the morning and evening may require higher PEPC activity than assumed in the simulations. Although predictions for total nitrogen investment into the thylakoids based on the evolutionary scenario are highly consistent with the measurements, the model overestimates the amount of cytochrome f by a factor of 2 (1.65 μmol m^−2^ instead of the measured 0.87 μmol m^−2^ for plants grown at 25°C, 1.43 μmol m^-2^ instead of 0.81 μmol m^−2^ at 35°C). However, the error of the measurements is uncertain, as no replicate measurements were performed for this parameter (Dwyer *et al.*, 2007). Discrepancies between model predictions and observations may also be in part due to error propagation from modeled amounts of chlorophyll and the photosystems. In each simulation, we optimized resource allocation for an environment that represents a static approximation to the dynamic environment a plant is facing. As diurnal and annual variations (which are no focus of this work) potentially show short-term trade-offs (Mori *et al.*, 2017; Reimers *et al.*, 2017), these might lead to a discrepancy between modeled and real evolutionary scenario. In particular, the difference between periodic and fluctuating conditions of the natural ancestral habitat on one hand, and the stable experimental growth conditions in audited growth chambers and the statically modeled evolutionary scenario on the other hand might have a strong effect.

In summary, we developed a general model of the complex photosynthetic apparatus, its resource requirements, and its interactions with environmental conditions. The presented modeling pipeline allows us to determine the extent of phenotypic plasticity and the relevance of different environmental conditions for photosynthetic organisms using C_3_, C_3_-C_4_ intermediate, and C_4_ metabolism. Applied to the physiological data from *Flaveria*, our work points to a strongly constrained phenotypic plasticity of C_4_ plants towards all considered environmental factors. This allows us to infer unique selective environments from plant performance and resource allocation data. More generally, our model provides a powerful tool to analyze the resource allocation of photosynthetic organisms and its dependence on environmental factors, allowing estimates for physiological and molecular parameters for which measurements are currently infeasible or impractical. This may prove to be of particular utility for systematically assessing the likely performance of crops in environments distinct from their natural habitats and for suggesting engineering targets in cases of limited phenotypic plasticity.

## Description

### Model overview

The nitrogen-dependent light-and enzyme-limited model allows us to calculate the environment-dependent net steady-state CO_2_ assimilation rate (*A*) of C_3_, C_4_, and all C_3_-C_4_ intermediate photosynthetic types. The model inputs are parameters defining the photosynthetic type and species-specific, invariable biochemical properties of the leaf to be modeled. Additionally, the input parameters comprise the following environmental factors: light intensity, leaf nitrogen level, temperature, and CO_2_ and O_2_ mesophyll partial pressures. We simulate a plant that is adapted to the input environment with respect to photosynthetic nitrogen and energy allocation. To this end, the nitrogen and energy allocation pattern that maximizes the net steady-state CO_2_ assimilation rate (*A*) is calculated via optimization, subject to the environmental and species-specific input parameters.

### Environmental factors and evolutionary parameters

We specify the environment in terms of the following factors: light intensity, leaf nitrogen level, temperature, and CO_2_ and O_2_ mesophyll partial pressures. The photosynthetic type is defined by six parameters: the Rubisco distribution between mesophyll and bundle sheath cells (β); the Rubisco kinetics, (specified through a single parameter, *k_ccat_* [μmol m^−2^ s^−1^], due to the known trade-off relationships between the kinetic parameters (Savir *et al.*, 2010)); the maximal C_4_ cycle activity (*V_pmax_*, [μmol m^−2^ s^−1^]); the fraction of glycine decarboxylated by the glycine decarboxylase complex in the bundle sheath cell that is derived from oxygenation by Rubisco in the mesophyll cell (*ξ*); the Michaelis constant of PEPC for bicarbonate (*K_p_*, [μbar]), and the bundle sheath cell conductance (*g_s_*, [μmol m^−2^ s^−1^]) (see Heckmann *et al.* (2013) for details). The values for the parameters are taken from the literature (see Supporting Information for details).

### Nitrogen allocation

To calculate the CO_2_ assimilation rate, we focus on the photosynthetic nitrogen pool (*N_ps_*, [μmol m^−2^]). In our model, *Nps* can be allocated across the following major pools of leaf photosynthetic nitrogen: the main enzyme of the Calvin-Benson cycle (*n_Etot_*), Rubisco; the main enzymes of the C_4_ cycle (*n_C4_*), PEPC and PPDK; and the thylakoids (*n_Jmax_*), which include the electron transport chains. *N_ps_* is calculated as a fraction of total leaf nitrogen (*N_t_*, [μmol m^−2^]) based on phenomenological observations according to Eqn 1, which comprises measured values for the investment into Rubisco, 12%, and the investment into the thylakoids (*n_fit_*, [fraction]) of C_3_ plants (Vogan & Sage, 2011; Vogan & Sage, 2012)*. n_fit_* represents a fit of the proportion of nitrogen invested into the thylakoids as a function of *N_t_*, based on the data of Vogan and Sage (2011).

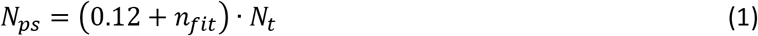

with

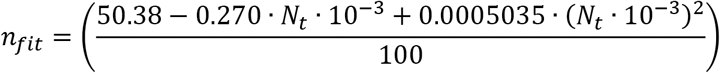

We assume a nitrogen investment into the photorespiratory enzymes of 13.8%, as suggested by Zhu *et al.* (2007) for a ‘typical’ C_3_ plant. To account for the reduced enzyme requirements of the photorespiratory cycle, we assume that *N_ps_* increases by 10% in plants that show sufficient C_4_ cycle activity; in our analyses, this applies to the C_3_-C_4_ intermediate, C_4_-like, and C_4_ species.

#### Nitrogen allocated to Rubisco

We only consider the nitrogen requirements of Rubisco in the Calvin-Benson cycle, as it accounts for the major nitrogen costs of this cycle (Evans & Seemann, 1989). The amount of catalytic sites of Rubisco (*Etot*, [μmol m^−2^]) is calculated from the invested nitrogen by Eqn 2, where *n_Etot_* represents the fraction of *Nps* invested into Rubisco:

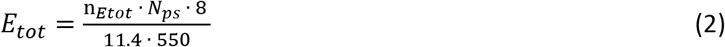

The parameters of this relationship are taken from Harrison *et al*. (2009).

#### Nitrogen allocated to enzymes of the C_4_ cycle

The nitrogen cost of C_4_ cycle enzymes is calculated from data on enzyme kinetics. The nitrogen requirements of the C_4_ cycle consider co-limitation of PEPC and PPDK, whose molecular weight (MW) and *kcat* are used to calculate the maximal rate of C_4_ cycle activity (Evans & von Caemmerer, 2000; Wang *et al.*, 2014). Eqn 3 represents the relationship between *V_pmax_* and nitrogen investment into the C_4_ enzymes (*n_C_*_4_*N_ps_*). MW* represents the nitrogen requirement of a catalytic site, assuming the nitrogen content is 16% (Makino *et al.*, 2003). Indices declare the considered enzyme.

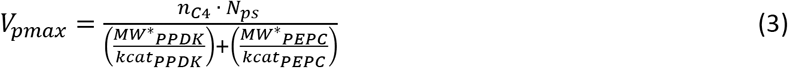

#### Nitrogen and the maximal electron transport rate

Nitrogen invested into the thylakoids (N_thy_ = N_t_ *n_thy_*, [μmol m^−2^]) is related to the maximal electron transport rate (*J_max_*, [μmol m^−2^ s^−1^]) via the amount of cytochrome f (cyt, [mmol/mol Chl]) and by considering photosystems I and II (PSI and PSII, [mmol/mol Chl]) as well as the light harvesting complexes (LHC, [mmol/mol Chl]). We use data from Ghannoum *et al*. (2005) for abundances of PSI and PSII to include phenomenological stoichiometry rules between LHC and the components of the electron transport chain (Eqns 4–8) and to relate *N_thy_* to the amount of cyt (Eqns 9–11). We assume that the chlorophyll content is shared between PSI, PSII, and LHC (Eqns 7 and 8). To be able to consider LET and CET, these complexes are split according to the proportion of LET (*p*) and CET (1 - *p*). Indices represent the considered pathway.

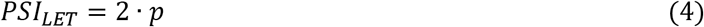

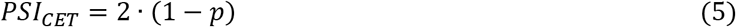

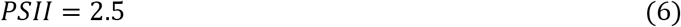

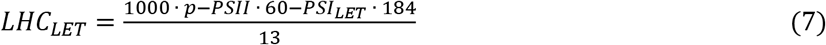

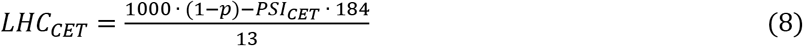

For the LET, *J_max_* is related to *N_thy_* as described in Eqns 9–12. *cyt_Jmax_* describes the relation of cyt to *Jmax* and was measured by Niinemets and Tenhunen (1997), who determined 156 (mmol e^-^)/(mmol cyt s) across various C_3_ species. Assuming 95% of LET in C_3_ plants, this leads to a capacity of 172 (mmol e^-^)/(mmol cyt s) for *cyt_Jmax_*.

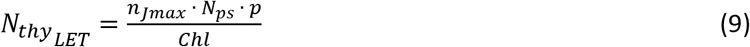

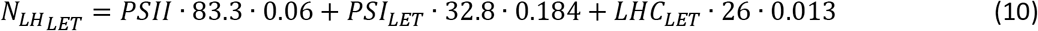

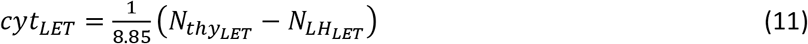

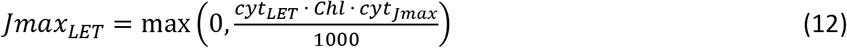

Chlorophyll content (*Chl*, [μmol m^−2^]) is calculated based on an empirical factor (Vogan & Sage, 2012) that relates the amount of nitrogen invested into thylakoids (*n_fit_ N_t_*, Eqn 1) to the amount of chlorophyll in C_3_ plants:

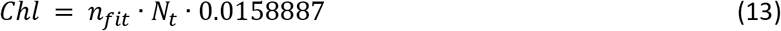

The response of chlorophyll content to leaf nitrogen does not differ significantly between different photosynthetic types in *Flaveria* (Vogan & Sage, 2011).

The derivation for the CET is analogous to the case of the LET (Eqns 14–17); additionally, the factor *Jmax_CL_* is required, which describes the scaling of *J_max_* with cyt for the CET:

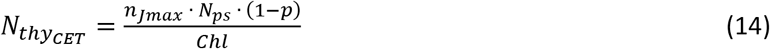

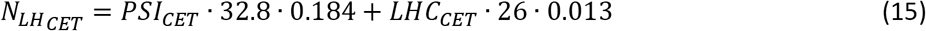

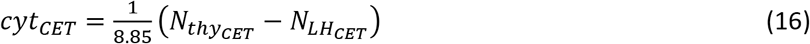

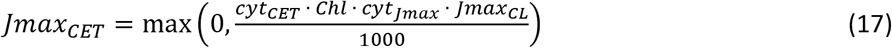

##### Optimization procedure

To find the maximal CO_2_ assimilation rate under the given environmental, physiological, and biochemical constraints, we optimize the allocation of photosynthetic nitrogen (assumed to depend only on total leaf nitrogen) into Rubisco, C_4_ cycle, LET, and CET through an augmented Lagrangian approach. The optimization is constrained to make sure that the results are biologically realistic, e.g., C_3_ species were not able to invest nitrogen into the C_4_ cycle (see Supporting Information for additional details).

The model and its optimization were implemented in the R environment (R Core Team, 2017), using the auglag-function of the package ‘nloptr’ (Johnson, see Supporting Information for details). The optimization algorithm can use various local solvers; we chose a derivative-free solver, ‘COBYLA’. We adapted the parameters of the auglag-function as follows: (1) *xtol_rel*=1×10^−100^, *i.e.*, we stop the optimization when all parameters changed by a proportion <1×10^−100^ in the last iteration; (2) *localtol*, the tolerance applied in the selected local solver, is set to 1×10^−100^; and (3) *maxeval*, the maximal number of optimization iterations, is set to 5×10^3^. To ensure robust retrieval of the global optimum, we perform a large number of optimizations starting from a wide range of initial values (see Supporting Information for details). The successful run resulting in the maximal CO_2_ assimilation rate is used.

### Modeling the effect of light

The relationship of the electron transport rate (*J_t_*, [μmol m^−2^ s^−1^]) and the absorbed light of a certain irradiance (*I*, [μmol m^−2^ s^−1^]) is presented in Eqns 18–20. *I* is related to *J_t_* by a widely accepted empirical hyperbolic function (Eqn 18), (von Caemmerer, 2000; Bernacchi *et al.*, 2003) that includes the following parameters: (1) *J_max_*, the maximum electron transport rate; (2) *Θ*, the convexity of the transition between the initial slope and the plateau of the hyperbola; (3) *α*, the leaf absorptance; (4) *f*, a correction factor accounting for the spectral quality of the light; and (5) *p,* the fraction of absorbed quanta that reaches PSI and PSII of LET (with (1 − *p*) reaching the CET). *I_abso_* is set to *I_LET_* and *I_CET_* dependent on the considered path of electron transport. The fraction of irradiance that is absorbed by the LET is shared equally between PSI and PSII (resulting in the factor 0.5 in Eqn 19), while the fraction of irradiance that is absorbed by the CET is assumed to reach PSI in full.

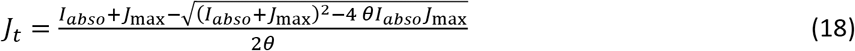

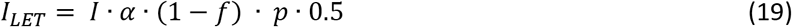

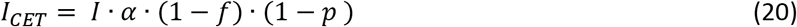

In our model it is assumed that the electron transport chain is the only source of ATP and NADPH and that both are used exclusively for CO_2_ fixation (von Caemmerer, 2000). As NADPH production results from LET, the amount of electrons is calculated using Eqns 18 and 19. The amount of electrons utilized for ATP production depends on both LET and CET (see below). There are multiple pathways of CET (Kramer & Evans, 2011); the model considers those pathways with an active Q-cycle and a ratio of two protons per electron. Note that Rubisco is assumed to be fully activated, independent of the irradiance (von Caemmerer, 2000).

The available energy needs to be partitioned between five pools: (1) the Calvin-Benson cycle (CBB) in the mesophyll; (2) the CBB in the bundle sheath; (3) the photorespiratory pathway (PR) in the mesophyll; (4) the PR in the bundle sheath cell; and (5) the C_4_ pathway. This means that the available energy is calculated in total and then partitioned (Kanai & Edwards, 1999) into *J_mp_*, *J_mc_,* and *J_s_*, the fractions invested into the C_4_ cycle, the CBB and the PR in the mesophyll, and the CBB and the PR in the bundle sheath cell, respectively. During optimization, the activity of each process is constrained by its allocated energy pool, *i.e.*, the energy allocation equals the relative energy allocation of the processes (see Supporting Information Method S1 for details).

The number of electrons transported to generate one molecule of ATP is unknown; for a discussion, see, e.g., Amthor (2010). We address these uncertainties by a factor that represents the ratio of electron transported per ATP in LET, which we set to *e_ATP_* = 4/3 in this work. In *Flaveria*, this ratio is supported by Siebke *et al.* (1997). The ATP and the NADPH requirements of the CBB, the PR, and the C_4_ cycle are based on the work of von Caemmerer (2000, see Supporting Information for equations). The energy requirements of the C_4_ cycle are adequate for the C_4_-subtypes that utilize NAD-malic enzyme or NADP-malic enzyme, whose ATP demand can be assumed to be equal. For the C_4_-subtype that utilizes PEP carboxykinase, the energetic costs are different and currently unclear (Kanai & Edwards, 1999; von Caemmerer, 2000).

### CO_2_ assimilation rate

A limitation in the production of both ATP and NADPH arises under light-limited conditions (von Caemmerer, 2000). The ATP-limited CO_2_ assimilation rate 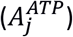 is calculated according to the light-limiting model of von Caemmerer (2000) (see Supporting Information for equations). The NADPH limitation is calculated analogously to the ATP-limited scenario (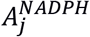, see Supporting Information).

The light-limited CO_2_ assimilation rate is then:

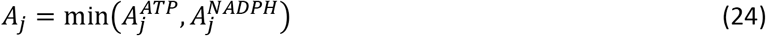

The model for the CO_2_ assimilation rate when the electron transport rate is not limiting (*A_c_*) is taken from Heckmann *et al.* (2013) and extended by a parameter representing the fraction of PSII activity in the bundle sheath cells, which affects O_2_ evolution. This parameter is set to *p*. In the whole model, each limitation is considered independently; the minimal CO_2_ assimilation rate determines the limiting process:

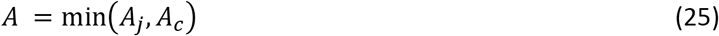

### Temperature-dependent model

Temperature affects the CO_2_ assimilation rate by changing the maximal activity of the C_4_ cycle, the carboxylation rate of Rubisco, and the electron transport rate. Temperature also affects the specificity of Rubisco as well as the Michaelis constants of Rubisco and PEPC. We model the temperature response by an extended Arrhenius function that describes two counteracting effects: rate increases with increasing temperature and enzyme inactivation through thermal instability (Massad *et al.*, 2007). We use parameters taken from literature or fitted to available data (see Supporting Information for the equation and a full list of parameters and their sources).

### Data used in the analyses

As the raw data of Vogan and Sage (2012) was not available, we extracted it from the corresponding figures using the Graph Grabber software provided by Quintessa Limited (Version 1.5.5). The measured curves consider the CO_2_ assimilation rate per intercellular CO_2_ concentration (*C_i_*). We assume that the mesophyll CO_2_ level is 85% of the *C_i_*.

For the detailed analysis of the C_4_ plants (Fig. 4), we used data published by Dwyer *et al.* (2007) for the CO_2_ assimilation rate at 25°C and 35°C, Rubisco catalytic sites, the PEPC activity, and the nitrogen investment into the thylakoids. As PEPC activity in *Flaveria* does not serve as a proxy for C_4_ cycle activity above values of around 130 μmol m^−2^ s^−1^ (Heckmann *et al.*, 2013), the maximal PEPC activity in C_4_ plants is set to 130 μmol m^−2^ s^−1^.

### Required nitrogen re-allocation (*δ_n_*)

Required nitrogen re-allocation (*δ_n_*, [fraction]) is defined as the total fraction of nitrogen that needs to be re-allocated between photosynthetic pools to optimally adjust photosynthesis from the evolutionary scenario 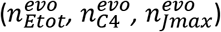 to a given growth environment 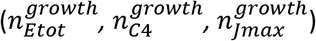:

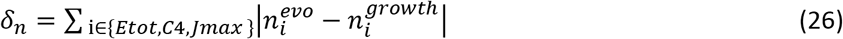

### Statistical information

The differences between adaptation scenarios are tested with the Wilcoxon rank sum test. Due to computational limitations, only a limited number of leaf nitrogen levels can be used to calculate the resource allocation for the data set of Vogan and Sage (2011) (Fig. 3). We considered 16 leaf nitrogen levels for the calculation of the resource allocation and CO_2_ assimilation rates. We inferred the CO_2_ assimilation rates required for the remaining leaf nitrogen levels from linear interpolation between the two closest leaf nitrogen levels. For the statistical analysis, the data of the modeled species, *F. pringlei* (C_3_), *F. floridana* (C_3_-C_4_), *F. palmeri* (C_4_-like), and *F. bidentis* (C_4_), was considered. All statistical analyses were conducted in R (R Core Team, 2017).

The difference of *δ_n_* for various photosynthetic types was tested by a sign test, applied to the data of Vogan and Sage (2011) (Supporting Information Table S5).

## Acknowledgements

We thank Prof. Rowan F. Sage and Prof. Susanne von Caemmerer for providing raw data from their publications. We further acknowledge financial support by the Deutsche Forschungsgemeinschaft (grants IRTG 1525 to DH, EXC 1028 to MJL and ES, CRC 680 and CRC 1310 to MJL). We would like to thank Bernhard O. Palsson and his group for hosting ES and for valuable discussions.

## Author Contributions

ES, MJL, and DH designed the research, interpreted the results, and wrote the paper. ES developed and implemented the model for nitrogen allocation and light reactions, and implemented the optimization procedure. DH developed and implemented the model for temperature responses. ES and DH conducted simulations and data analysis.

## Supporting Information

Additional supporting information may be found in the online version of this article.

**Methods S1** Details about the optimization procedure of resource allocation

**Methods S2** Equations of the energetic costs

**Methods S3** Equations of the light-limited CO_2_ assimilation rate

**Methods S4** Details about the temperature-dependent model

**Methods S5** Sensitivity analysis

**Table S1** *Flaveria* parametrization.

**Table S2** Lower and upper bounds for the model parameters subject to numerical optimization.

**Table S3** The parameters of the temperature-dependent model.

**Table S4** Required nitrogen re-allocation (*δ_n_*) for *F. bidentis* (C_4_) grown at different temperatures.

**Table S5** Required nitrogen re-allocation (*δ_n_*) for different on leaf nitrogen level for various *Flaveria* species.

**Table S6** Required nitrogen re-allocation (*δ_n_*) for various *Flaveria* species grown at current or low CO_2_ level.

**Table S7** The modeled and measured data of chlorophyll and PSII of *F. bidentis* (C_4_).

**Table S8** Distribution parameters used to generate the random parameter sets for the sensitivity.

**Fig. S1** Sensitivity analysis.

**Fig. S2** A-C_i_ curve measured at 40°C using plants grown at the current CO_2_ level.

**Fig. S3** A-C_i_ curve measured at 30°C using plants grown at the low CO_2_ level.

**Fig. S4** A-C_i_ curve measured at 40°C using plants grown at the low CO_2_ level.

**Fig. S5** A-Temperature curve using plants grown at the low CO_2_ level

**Fig. S6** Discrepancy between measured and modeled results of *F. robusta* (C_3_) across diverse environments assuming no phosphate-limitation.

**Fig. S7** Discrepancy between measured and modeled results of *F. robusta* (C_3_) across diverse environments assuming phosphate-limitation.

## References

Amthor JS. 2010. From sunlight to phytomass: on the potential efficiency of converting solar radiation to phyto-energy. New Phytologist 188(4): 939-959.

Atkinson D. 1969. Limitation of metabolite concentrations and the conservation of solvent capacity in the living cell. Current topics in cellular regulation 1: 29-43.

Baudouin-Cornu P, Surdin-Kerjan Y, Marliere P, Thomas D. 2001. Molecular evolution of protein atomic composition. Science 293(5528): 297-300.

Beg QK, Vazquez A, Ernst J, de Menezes MA, Bar-Joseph Z, Barabasi AL, Oltvai ZN. 2007. Intracellular crowding defines the mode and sequence of substrate uptake by Escherichia coli and constrains its metabolic activity. Proceedings of the National Academy of Sciences of the United States of America 104(31): 12663-12668.

Bernacchi CJ, Pimentel C, Long SP. 2003. In vivo temperature response functions of parameters required to model RuBP-limited photosynthesis. Plant Cell and Environment 26(9): 1419-1430.

Berry JA, Farquhar GD 1978. The CO_2_ concentrating function of C_4_ photosynthesis: a biochemical model. Proceedings of the Fourth International Congress on Photosynthesis. Biochemical Society, London. 119-131.

Christin PA, Sage TL, Edwards EJ, Ogburn RM, Khoshravesh R, Sage RF. 2011. Complex Evolutionary Transitions and the Significance of C_3_-C_4_ Intermediate Forms of Photosynthesis in Molluginaceae. Evolution 65(3): 643-660.

de Oliveira Dal’Molin CG, Quek LE, Palfreyman RW, Brumbley SM, Nielsen LK. 2010. AraGEM, a genome-scale reconstruction of the primary metabolic network in Arabidopsis. Plant Physiology 152: 579.

Dourado H, Maurino VG, Lercher MJ. 2017. Enzymes And Substrates Are Balanced At Minimal Combined Mass Concentration In Vivo. bioRxiv.

Dwyer SA, Ghannoum O, Nicotra A, Von Caemmerer S. 2007. High temperature acclimation of C_4_ photosynthesis is linked to changes in photosynthetic biochemistry. Plant Cell and Environment 30(1): 53-66.

Ellis RJ. 1979. Most Abundant Protein in the World. Trends in Biochemical Sciences 4(11): 241-244.

Evans JR, Seemann JR. 1989. The allocation of protein nitrogen in the photosynthetic apparatus: costs, consequences, and control. Photosynthesis: 183-205.

Evans JR, von Caemmerer S 2000. Would C_4_ rice produce more biomass than C_3_ rice? In: Sheehy JE, Mitchell PL, Hardy B eds. Redesigning rice photosynthesis to increase yield: Elsevier, 53-71.

Farquhar GD, Caemmerer S, Berry JA. 1980. A biochemical model of photosynthetic CO_2_ assimilation in leaves of C_3_ species. Planta 149: 78-90.

Friend AD. 1991. Use of a Model of Photosynthesis and Leaf Microenvironment to Predict Optimal Stomatal Conductance and Leaf Nitrogen Partitioning. Plant Cell and Environment 14(9): 895-905.

Gerhart LM, Ward JK. 2010. Plant responses to low CO_2_ of the past. New Phytologist 188(3): 674-695.

Ghannoum O, Evans JR, Chow WS, Andrews TJ, Conroy JP, von Caemmerer S. 2005. Faster rubisco is the key to superior nitrogen-use efficiency in NADP-malic enzyme relative to NAD-malic enzyme C_4_ grasses. Plant Physiology 137(2): 638-650.

Harrison MT, Edwards EJ, Farquhar GD, Nicotra AB, Evans JR. 2009. Nitrogen in cell walls of sclerophyllous leaves accounts for little of the variation in photosynthetic nitrogen-use efficiency. Plant Cell and Environment 32(3): 259-270.

Heckmann D, Schulze S, Denton A, Gowik U, Westhoff P, Weber A PM, Lercher Martin J. 2013. Predicting C_4_ Photosynthesis Evolution: Modular, Individually Adaptive Steps on a Mount Fuji Fitness Landscape. Cell 153(7): 1579-1588.

Ibarra RU, Edwards JS, Palsson BO. 2002. Escherichia coli K-12 undergoes adaptive evolution to achieve in silico predicted optimal growth. Nature 420(6912): 186-189.

Johnson SG The NLopt nonlinear-optimization package.

Kanai R, Edwards GE 1999. The biochemistry of C_4_ photosynthesis. In: Sage RF, Monson RK eds. C4 plant biology: Academic press, Toronto, ON, Canada, 49-87.

Kramer DM, Evans JR. 2011. The Importance of Energy Balance in Improving Photosynthetic Productivity. Plant Physiology 155(1): 70-78.

Maire V, Martre P, Kattge J, Gastal F, Esser G, Fontaine S, Soussana JF. 2012. The Coordination of Leaf Photosynthesis Links C and N Fluxes in C_3_ Plant Species. PLoS ONE 7(6).

Makino A, Sakuma H, Sudo E, Mae T. 2003. Differences between maize and rice in N-use efficiency for photosynthesis and protein allocation. Plant and Cell Physiology 44(9): 952-956.

Malhi SS, Grant CA, Johnston AM, Gill KS. 2001. Nitrogen fertilization management for no-till cereal production in the Canadian Great Plains: a review. Soil & Tillage Research 60(3-4): 101-122.

Mallmann J, Heckmann D, Bräutigam A, Lercher MJ, Weber AP, Westhoff P, Gowik U. 2014. The role of photorespiration during the evolution of C_4_ photosynthesis in the genus *Flaveria*. eLife 10.7554/eLife.02478.

Massad RS, Tuzet A, Bethenod O. 2007. The effect of temperature on C_4_-type leaf photosynthesis parameters. Plant Cell and Environment 30(9): 1191-1204.

Maurino VG, Peterhansel C. 2010. Photorespiration: current status and approaches for metabolic engineering. Current Opinion in Plant Biology 13(3): 249-256.

McKown AD, Moncalvo J-M, Dengler NG. 2005. Phylogeny of *Flaveria* (Asteraceae) and inference of C_4_ photosynthesis evolution. American Journal of Botany 92: 1911-1928.

Mori M, Schink S, Erickson DW, Gerland U, Hwa T. 2017. Quantifying the benefit of a proteome reserve in fluctuating environments. Nature Communications 8.

Munekage YN, Taniguchi YY. 2016. Promotion of Cyclic Electron Transport Around Photosystem I with the Development of C_4_ Photosynthesis. Plant and Cell Physiology 57(5): 897-903.

Niinemets U, Tenhunen JD. 1997. A model separating leaf structural and physiological effects on carbon gain along light gradients for the shade-tolerant species *Acer saccharum*. Plant Cell and Environment 20(7): 845-866.

Oberhardt MA, Palsson BO, Papin JA. 2009. Applications of genome-scale metabolic reconstructions. Molecular Systems Biology 5.

Powell AM. 1978. Systematics of *Flaveria* (Flaveriinae-Asteraceae). Annals of the Missouri Botanical Garden: 590-636.

R Core Team 2017. R: A Language and Environment for Statistical Computing: R Foundation for Statistical Computing.

Reimers AM, Knoop H, Bockmayr A, Steuer R. 2017. Cellular trade-offs and optimal resource allocation during cyanobacterial diurnal growth. Proc Natl Acad Sci U S A.

Sage RF. 2004. The evolution of C_4_ photosynthesis. New Phytologist 161: 341-370.

Sage RF, Cowling SA. 1999. Implications of stress in low CO_2_ atmospheres of the past: are today’s plants too conservative for a high CO_2_ world. Carbon dioxide and environmental stress: 289-308.

Sage RF, McKown AD. 2006. Is C_4_ photosynthesis less phenotypically plastic than C_3_ photosynthesis? Journal of Experimental Botany 57(2): 303-317.

Savir Y, Noor E, Milo R, Tlusty T. 2010. Cross-species analysis traces adaptation of Rubisco toward optimality in a low-dimensional landscape. Proceedings of the National Academy of Sciences 107(8): 3475-3480.

Siebke K, vonCaemmerer S, Badger M, Furbank RT. 1997. Expressing an RbcS antisense gene in transgenic Flaveria bidentis leads to an increased quantum requirement for CO2 fixed in photosystems I and II. Plant Physiology 115(3): 1163-1174.

Vance CP. 2001. Symbiotic nitrogen fixation and phosphorus acquisition. Plant nutrition in a world of declining renewable resources. Plant Physiology 127(2): 390-397.

Vogan PJ, Sage RF. 2011. Water-use efficiency and nitrogen-use efficiency of C_3_-C_4_ intermediate species of *Flaveria* Juss. (Asteraceae). Plant, Cell & Environment 34: 1415-1430.

Vogan PJ, Sage RF. 2012. Effects of low atmospheric CO_2_ and elevated temperature during growth on the gas exchange responses of C_3_, C_3_-C_4_ intermediate, and C_4_ species from three evolutionary lineages of C_4_ photosynthesis. Oecologia 169(2): 341-352.

von Caemmerer S. 1989. A model of photosynthetic CO_2_ assimilation and carbon-isotope discrimination in leaves of certain C_3_−C_4_ intermediates. Planta 178(4): 463-474.

von Caemmerer S. 2000. Biochemical models of leaf photosynthesis. Collingwood, Australia: Csiro Publishing.

Wang Y, Long SP, Zhu XG. 2014. Elements Required for an Efficient NADP-Malic Enzyme Type C_4_ Photosynthesis. Plant Physiology 164(4): 2231-2246.

Zhu X-G, de Sturler E, Long SP. 2007. Optimizing the distribution of resources between enzymes of carbon metabolism can dramatically increase photosynthetic rate: a numerical simulation using an evolutionary algorithm. Plant Physiol 145: 513-526.

